# Altered collective mitochondrial dynamics in an *Arabidopsis msh1* mutant compromising organelle DNA maintenance

**DOI:** 10.1101/2021.10.22.465420

**Authors:** Joanna M. Chustecki, Ross D. Etherington, Daniel J. Gibbs, Iain G. Johnston

## Abstract

Mitochondria form highly dynamic populations in the cells of plants (and all eukaryotes). The characteristics of this collective behaviour, and how it is influenced by nuclear features, remain to be fully elucidated. Here, we use a recently-developed quantitative approach to reveal and analyse the physical and collective “social” dynamics of mitochondria in an *Arabidopsis msh1* mutant where organelle DNA maintenance machinery is compromised. We use a newly-created line combining the *msh1* mutant with mitochondrially-targeted GFP, and characterise mitochondrial dynamics with a combination of single-cell timelapse microscopy, computational tracking and network analysis. The collective physical behaviour of *msh1* mitochondria is altered from wildtype in several ways: mitochondria become less evenly spread, and networks of inter-mitochondrial encounters become more connected with greater potential efficiency for inter-organelle exchange. We find that these changes are similar to those observed in *friendly*, where mitochondrial dynamics are altered by a physical perturbation, suggesting that this shift to higher connectivity may reflect a general response to mitochondrial challenges.

## Introduction

Mitochondria are key bioenergetic compartments of the eukaryotic cell. Within plant cells, hundreds of mitochondria exist, largely as individual organelles -- contrasting with the reticulated network form often seen in yeast and mammalian cells (Logan, 2006b; Johnston, 2019). These cellular populations are highly dynamic (Logan, 2010), interacting with each other and other organelles (Islam, Niwa and Takagi, 2009; Jaipargas *et al*., 2015; Shai, Schuldiner and Zalckvar, 2016; Barton *et al*., 2018; Krupinska *et al*., 2020; Chustecki *et al*., 2021).

Recent work suggested that the collective cellular dynamics of plant mitochondria can resolve a tension between mitochondrial proximity and spacing (Chustecki *et al*., 2021). Mitochondria need to be physically proximal to allow membrane fusion and mixing of contents including mitochondrial DNA (mtDNA) (Arimura *et al*., 2004; Sheahan, McCurdy and Rose, 2005; Rose, 2021). In addition to this exchange, mitochondrial proximity facilitates metabolic exchange and mitochondrial quality control, a process reliant on cycles of fission and fusion, key for maintaining a healthy chondriome (Jones, 1986; Karbowski and Youle, 2003; Arimura *et al*., 2004; Logan, 2006a; Takanashi *et al*., 2006; Twig *et al*., 2008; Liu *et al*., 2009; Figge *et al*., 2012; Shutt and McBride, 2013; Agrawal, Pekkurnaz and Koslover, 2018). There are also many other functional implications of inter-mitochondrial proximity including influence on membrane potential (Santo-Domingo *et al*., 2013), cristae alignment (Picard *et al*., 2015), and calcium waves (Ichas, Jouaville and Mazat, 1997). However, there are also benefits to mitochondria remaining physically spaced, with benefits for energy demand, inter-organellar colocalisation, and the regulation of metabolic demands (Chen and Chan, 2006; Seguí-Simarro and Staehelin, 2009; Bauwe, Hagemann and Fernie, 2010; Sage, Sage and Kocacinar, 2012; Liesa and Shirihai, 2013; Spillane *et al*., 2013; Shai, Schuldiner and Zalckvar, 2016; Yu *et al*., 2016; Schuler *et al*., 2017; Yu and Pekkurnaz, 2018). The mitochondrial population thus faces a hypothesised tension between maintaining even spacing of mitochondria and supporting inter-mitochondrial encounters.

Chustecki *et al*. (2021) explored this tradeoff between even spacing and supporting encounters by characterising the ‘social networks’ of the dynamic cellular population, allowing the characterisation of connectivity across the chondriome -- the whole population of mitochondria in a cell (Logan, 2010). Physical and network analysis revealed that wildtype *Arabidopsis* uses mitochondrial dynamics to resolve this tension, with mitochondrial motion allowing transient encounters between organelles -- and facilitating efficient exchange through the population -- while also retaining physical spacing. The development of this approach allows targeted, quantitative questions to be asked about how collective mitochondrial behaviour responds to different situations.

Here, we pursue this target by investigating the collective behaviour of mitochondria in the *msh1* mutant, where MutS HOMOLOGUE 1 (MSH1), responsible for recombination surveillance of organellar genomes (Martínez-Zapater *et al*., 1992; Abdelnoor *et al*., 2003, 2006; Shedge *et al*., 2007; Arrieta-Montiel *et al*., 2009; Davila *et al*., 2011; Wu *et al*., 2020), is compromised. Disruption of mitochondrial-localised MSH1 leads to an increase in single nucleotide variants and insertion-deletion mutations in mtDNA (Wu *et al*., 2020). In some plants, MSH1 disruption can also lead to substoichiometric shifting in the mitochondrial genome (Martínez-Zapater *et al*., 1992; Sakamoto *et al*., 1996; Abdelnoor *et al*., 2003). Although the full molecular mechanism of MSH1 action on the mitochondrial genome is still not fully characterised (Fukui *et al*., 2018; Wu *et al*., 2020), multiple studies support the model of MSH1 influencing double strand break repair (Davila *et al*., 2011; Christensen, 2014; Wu *et al*., 2020). *msh1* does not, however, exclusively affect mtDNA: chloroplast DNA maintenance is also compromised, and other effects, including metabolic influences of the resulting organelle dysfunction and even epigenetic changes likely also contribute to the phenotype (Xu *et al*., 2011, 2012; Virdi *et al*., 2015; Shao *et al*., 2017).

Disruption of MSH1 thus provides genetic challenges to the mtDNA and plastid DNA (ptDNA) populations, as well as resultant metabolic and other stresses. We set out to investigate whether these stresses had the effect of changing the collective cellular behaviour of mitochondria. As described below, we explored this question by using single-cell microscopy, computational analysis and network science approaches to characterise and analyse mitochondrial behaviour in *msh1* compared to wildtype *Arabidopsis* and other mutants.

## Results

### Construction, genotyping and phenotyping of mtGFP-*msh1*

To allow the visualisation of mitochondrial dynamics in the *msh1* mutant, we created mtGFP-*msh1*, combining the transgenic mtGFP line where GFP is localised to mitochondria (from an original line kindly provided by Prof David Logan (Logan and Leaver, 2000)) with a mutant line where MSH1, an organelle genome maintenance factor, is perturbed by a premature stop codon-caused by a single nucleotide polymorphism (SNP) (Abdelnoor *et al*., 2003; see Methods for more details). We verified the crossed line using dCAPs genotyping for the SNP and rosette phenotyping for characteristic variegation in the *msh1* line (SI Fig. 1), where in contrast to both wildtype mtGFP and Col-0, mtGFP-*msh1* retained the expected variegated and low growth phenotype of the *msh1* mutant (SI Fig. 2A,B,C). The candidate line at F3 showed the presence of the SNP (SI Fig. 1A), as well as resistance to Kanamycin, demonstrating presence of the mtGFP transgene (Logan and Leaver, 2000). Sequencing of the F3 candidate line confirmed the presence of the SNP in the region encoding *MSH1* (SI Fig. 3). Sequencing of three F4 candidate line offspring also showed the presence of the SNP, validating the genetic makeup of the mtGFP-*msh1* mutant.

### *msh1* alters physical dynamics of mitochondria

Following the creation of mtGFP-*msh1*, we used confocal microscopy to characterise mitochondrial dynamics in single hypocotyl cells of 4-5 day seedlings in this mutant, and compared these dynamics to the mtGFP transgenic line, representing wildtype mitochondrial motion. This imaging approach followed the protocol from (Chustecki *et al*., 2021). Briefly, we recorded timelapse videos of mitochondrial motion in single cells, and computationally identified trajectories of individual mitochondria using TrackMate (Tinevez *et al*., 2017). From these trajectories we can analyse individual and collective behaviour of mitochondria, including speeds, colocalisations, and many more statistics (Chustecki *et al*., 2021). Fig. 1 illustrates the process of tracking fluorescent mitochondria over time, in representative mtGFP (Fig. 1Ai) and mtGFP-*msh1* (Fig. 1Bi) single cells. Generally and qualitatively, as with wildtype mtGFP mitochondrial motion, mtGFP-*msh1* mitochondria showed a mixture of diffusive and ballistic motion, with some organelles remaining static, and others moving swiftly across the cell. These organelles also colocalise with one another, and occasionally colocalise with chloroplasts (Supp Video 1).

**Figure 1:**
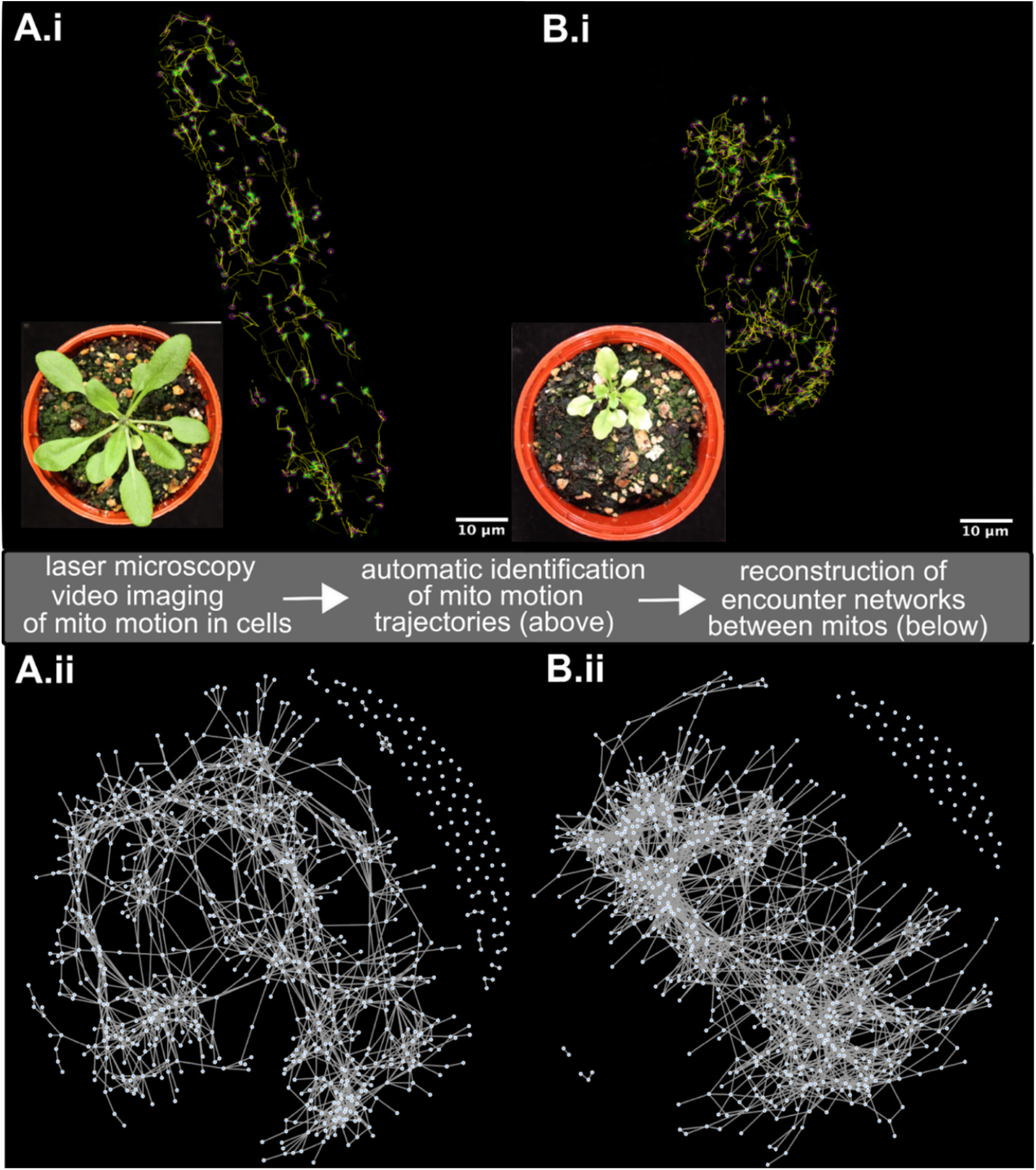
Characterising the “social networks” of plant mitochondria in mtGFP. **(A) and mtGFP-*msh1* (B)**. Top panels (i) illustrate the tracking process of (green) fluorescent mitochondria in single hypocotyl cells from seedlings, using Trackmate (Tinevez *et al*., 2017). Mitochondria are automatically identified (pink spots, diameter 1μm), and computed tracks over time are shown (for clarity, only 10 local frames are shown (yellow)). Insets show whole-plant phenotypes of the two lines at later development. Bottom panels (ii) show the networks of mitochondrial encounters corresponding to the single-cell dynamics (nodes are mitochondria, edges are encounters), built up over a time window of observation (here 233 seconds).

We found that mitochondria in mtGFP-*msh1* on average were less evenly spread and were physically associated for longer times in hypocotyl cells (Fig. 2). Mean inter-mitochondrial distance, reporting the average distance (in microns) to the nearest physical neighbour in the cell, was lower in mtGFP-*msh1*, reflecting a less evenly-spread population (Fig. 2A). The median speed of individual mitochondria in mtGFP-*msh1* was also lower, although differences between the lines did not cross a significance threshold when we used a conservative non-parametric test (Fig. 2B). Colocalisation time, reporting the time over which two mitochondria are within a threshold of each other, was higher in mtGFP-*msh1* (Fig. 2C). Cell sizes were similar across all lines (SI Fig. 5), suggesting that these physical differences are intrinsic properties of the mitochondrial population and not a result of altered cellular morphology.

**Figure 2:**
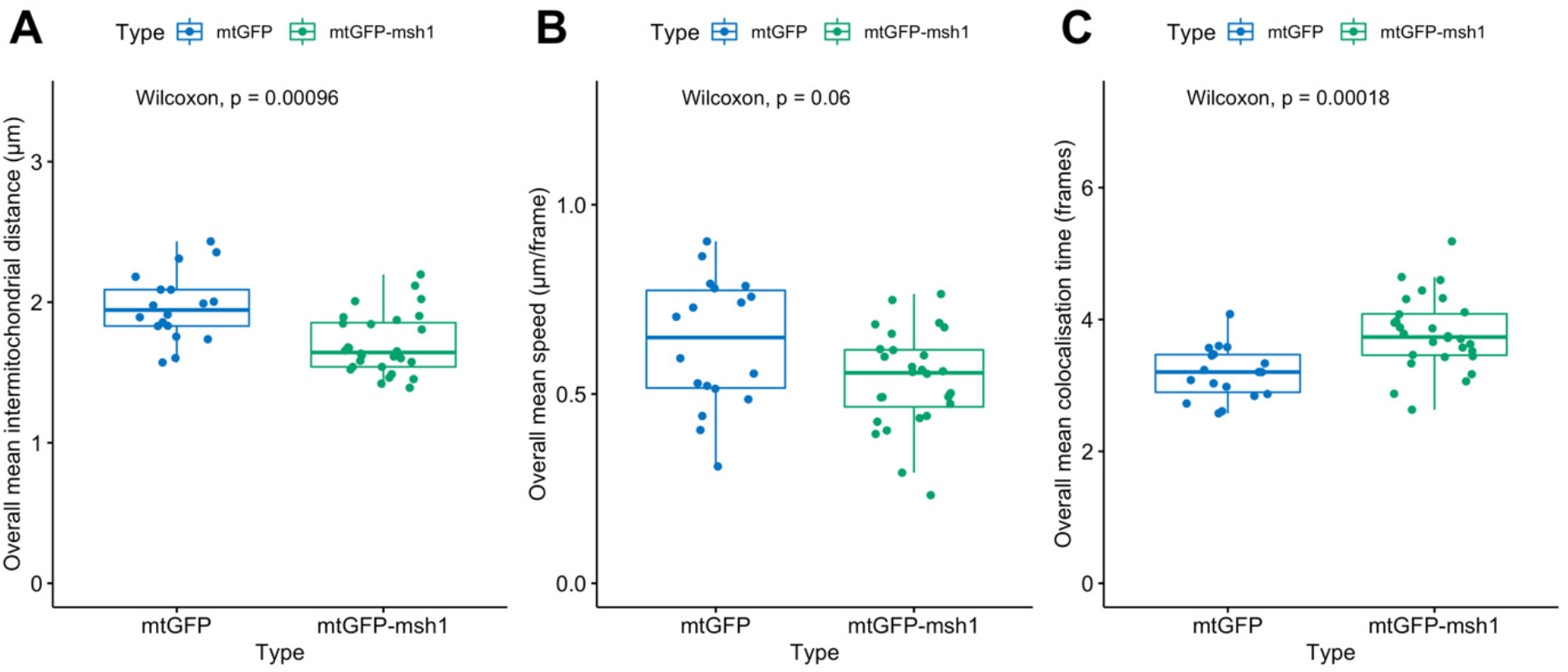
Physical summary statistics differ between mtGFP and mtGFP-*msh1*. Each point represents a summary statistic for one cell (mtGFP *n*= 18, mtGFP-*msh1 n*= 28). P-values represents outcome of the Wilcoxon rank sum test across both genotypes, without multiple hypothesis correction. Boxplots represent the median and 25th/75th percentile, with whiskers showing the smallest/largest value within 1.5x the interquartile range. Each individual point gives the mean statistic across an entire video, corresponding to 233 seconds of video time.

### Alterations in physical dynamics of *msh1* affect social dynamics

To explore whether this decrease in spacing is accompanied by an increase in inter-mitochondrial connectivity, we next characterised the “encounter networks” of mitochondria, defined as the set of colocalisations between pairs of mitochondria that occur within a given timeframe (see Methods, Fig. 1Aii, Bii, SI Fig. 4). Akin to social networks, describing social interactions between individuals in a population, these encounter networks shape the potential for exchange of contents across the mitochondrial population (Chustecki *et al*., 2021).

Salient features of these encounter networks for collective mitochondrial behaviour are the degree distribution (the number of different mitochondria each mitochondrion encounters); the diameter of the network (the length in edges of the longest direct route across the network) and the network efficiency. This final quantity is the average of the reciprocal lengths of the shortest paths between each pair of mitochondria in the network. If all pairs of mitochondria are connected by short paths (facilitating exchange through the network), reciprocal lengths, and network efficiency, are high. If some pairs are connected only by long paths, or are disconnected, reciprocal lengths and efficiency are low and information exchange is more challenging.

We found that the encounter networks of mtGFP-*msh1* had higher mean degree and higher efficiency than the mtGFP single mutant (representative of wildtype mitochondrial networks) (Fig. 3A,B). Mitochondria in the *msh1* mutant are thus more directly connected through encounters, facilitating easier exchange of contents. Network diameter is also shorter across mtGFP-*msh1* networks, again suggesting increased organelle connectivity; but we note the significant difference was not retained after multiple hypothesis testing (Fig. 3C). The size of networks, quantified either by node or edge number, remained similar between mtGFP and mtGFP-*msh1* over time (SI Fig. 6). There was no significant difference across values for betweenness centrality, an average of the number of shortest paths crossing each node in the network (Fig. 3D).

**Figure 3:**
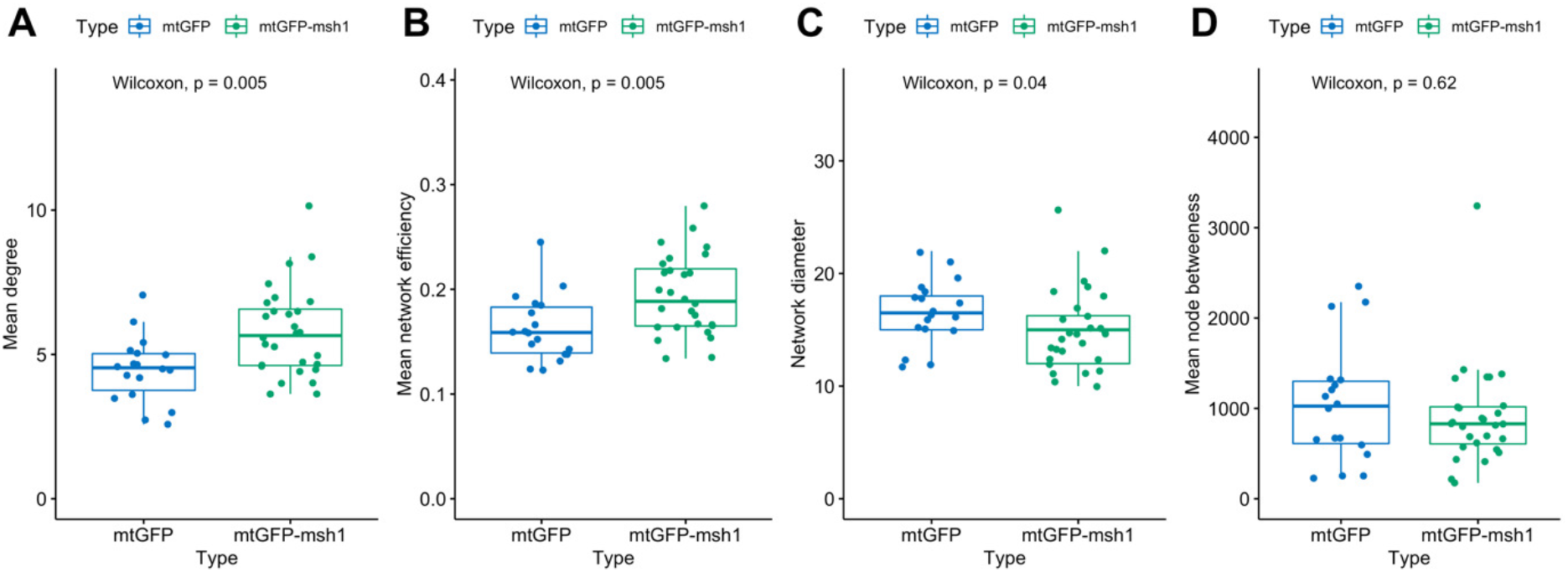
Social summary statistics differ between mtGFP and mtGFP-*msh1*. Each point represents a summary statistic for one cell (mtGFP *n*= 18, mtGFP-*msh1 n*= 28). P-values represents outcome of the Wilcoxon rank sum test across both genotypes, without multiple hypothesis correction. Boxplots represent the median and 25th/75th percentile, with whiskers showing the smallest/largest value within 1.5x the interquartile range. Each individual point is from a network corresponding to an observed time window of 233 seconds.

These network statistics are time-dependent, because networks build up over time as more encounters between individuals occur. As seen in SI Fig. 7, *msh1* differences in degree value remain across observation time windows, with network efficiency differences significant at later frames (SI Fig. 7A,B, Fig. 3A,B), when networks have built up with more encounters. Network diameter relationships across the lines do not change over time, but betweenness centrality is significantly different for early comparisons between lines, but not at later frames (SI Fig. 7C,D, Fig. 3C,D). This could be a consequence of the topology of smaller networks, before so many encounters and connections between smaller cliques of mitochondria are formed.

### The collective dynamic response to *msh1* resembles the response to *friendly*

We next asked whether the altered mitochondrial behaviour in the face of the *msh1* perturbation shared similarities with altered behaviour under a physical perturbation to mitochondrial dynamics. To this end, we characterised an mtGFP-*friendly* mutant within which the fusion of these organelles is perturbed (El Zawily *et al*., 2014), increasing the association time between individuals, and posing a transient challenge to the social connectivity and physical spread trade-off as shown in (Chustecki *et al*., 2021). Recent work has illuminated the colocalization of FRIENDLY to depolarised mitochondria as an essential part of the mitophagy pathway (Ma *et al*., 2021) -- its perturbation results in reduced mitochondrial fusion, increased mitochondrial clustering, and a wide range of metabolic issues (El Zawily *et al*., 2014; Ma *et al*., 2021). This mutant has a pronounced growth phenotype, though more limited than *msh1* (SI Fig. 2D).

To explore the relationship between changes in mitochondrial behaviour due to physical and genetic challenges, we compared mitochondrial behaviour in mtGFP, mtGFP-*msh1*, and mtGFP-*friendly*. Strikingly, the physical and social statistics observed in mtGFP-*msh1* and mtGFP-*friendly* lines are remarkably similar, with no statistically detectable differences between these genotypes. Of course, an absence of statistical significance does not imply the absence of an effect, but the observed magnitudes of the statistics and our moderate sample sizes (*n*=28 for mtGFP-*msh1, n*=19 for mtGFP-*friendly*) suggest that the behaviours are indeed rather similar (Fig. 4). There was a slightly lower inter-mitochondrial distance alongside an increased degree and network efficiency within mtGFP-*msh1* -- suggesting a marginally more pronounced shift towards connectivity -- although these observations did not meet a statistical significance threshold for a nonparameteric comparison (Fig. 4 A,D,E). Both mutant genotypes show a significantly decreased inter-mitochondrial distance, and increased colocalization time and degree, when compared to wildtype mtGFP (Fig. 4 A,C,D).

**Figure 4:**
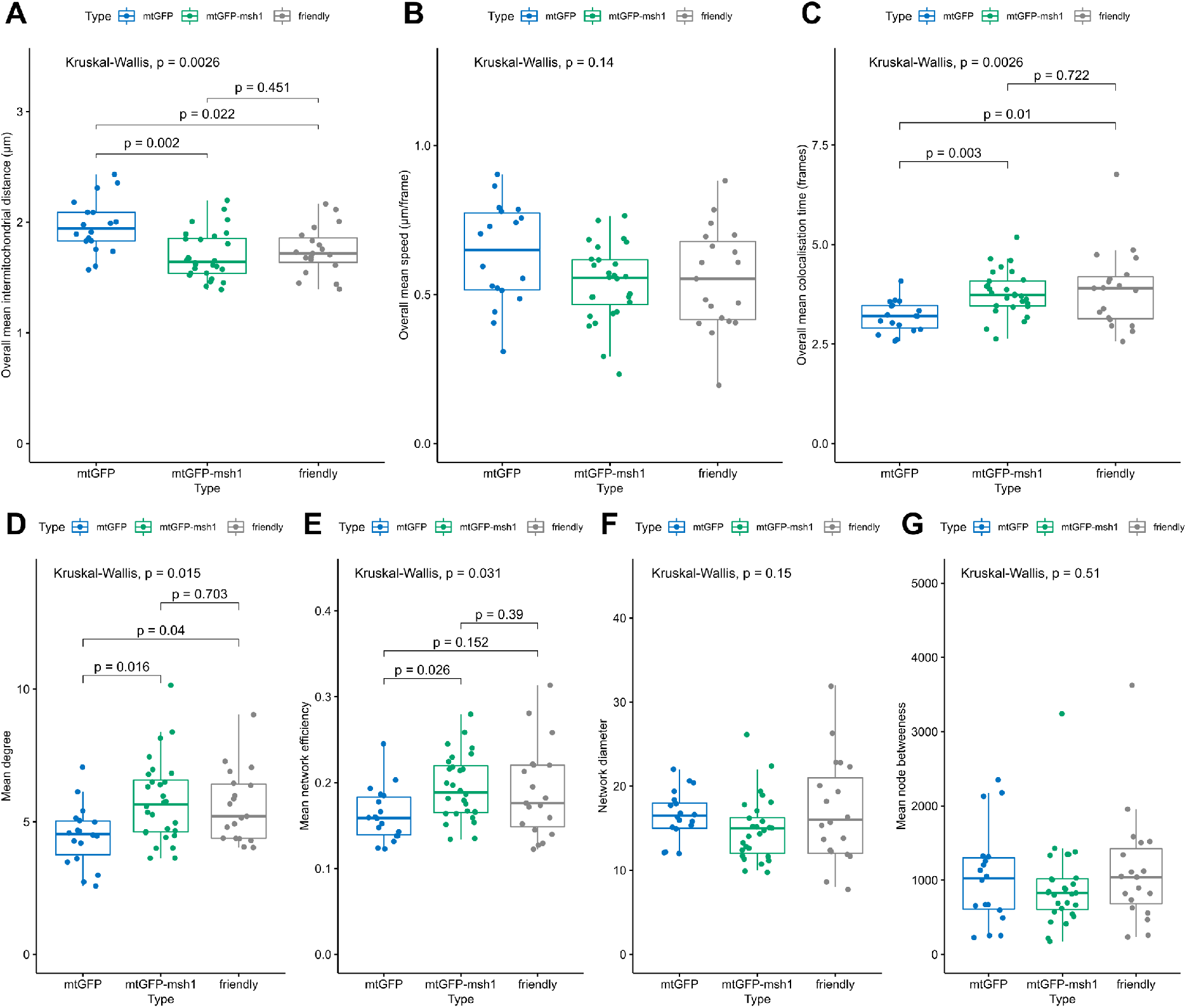
Physical and social summary statistics compared across mtGFP, mtGFP-*msh1* and *friendly*. Each point represents a summary statistic for one cell (mtGFP *n*= 18, mtGFP-*msh1 n*= 28, *friendly n* = 19). P-values represent Kruskal Wallis test outcomes across all three genotypes, and pairwise p-values are false discovery rate adjusted outcomes of a post-hoc Dunn test, without multiple hypothesis correction across statistics. Boxplots represent the median and 25th/75th percentile, with whiskers showing the smallest/largest value within 1.5x the interquartile range. Each physical datapoint (A-C) is a mean across a 233 second time window, and each social datapoint (D-G) is from a network corresponding to a time window of 233 seconds.

Previous work (Chustecki *et al*., 2021) found that the difference between mtGFP-*friendly* and wildtype behaviour diminished over time: initially rather cliquey, the *friendly* networks became more globally connected over time as itinerant mitochondria formed social bridges between cliques. Our statistical analysis here supports this picture for mean degree in both *friendly* and *msh1* (SI Fig. 7A; Fig. 4D) while revealing a more nuanced picture for other network statistics. In particular, network efficiency differences between the mutants and wildtype do not diminish over time to the same extent (SI Fig. 7B; Fig. 4E), suggesting that the global changes in collective behaviour are maintained robustly over time despite similarities in local behaviour. Overall, both the magnitudes and time behaviour of collective dynamic changes were quantitatively similar in *friendly* and *msh1*, supporting the comparable influences of the two perturbations.

## Discussion

Mitochondria across eukaryotes are strikingly dynamic. In some cases, including the delivery of ATP to synapses in neurons (Hollenbeck and Saxton, 2005; Mironov, 2007; MacAskill, Atkin and Kittler, 2010) and fit mitochondria to growing buds in yeast (Fehrenbacher *et al*., 2004; Pernice *et al*., 2018), the reasons for this motion are largely explained. In many other cases, the advantages and disadvantages of the rich dynamics of mitochondria remain unclear. Here we have demonstrated that two perturbations to nuclear-encoded machinery, *msh1* and *friendly*, influence the collective dynamics of plant mitochondria in a similar way: trading reduced spacing for increased connectivity. The quantitative similarity between the two responses suggests that this shift may reflect a more general response of plant cells to organelle stress (Fig. 5).

**Figure 5:**
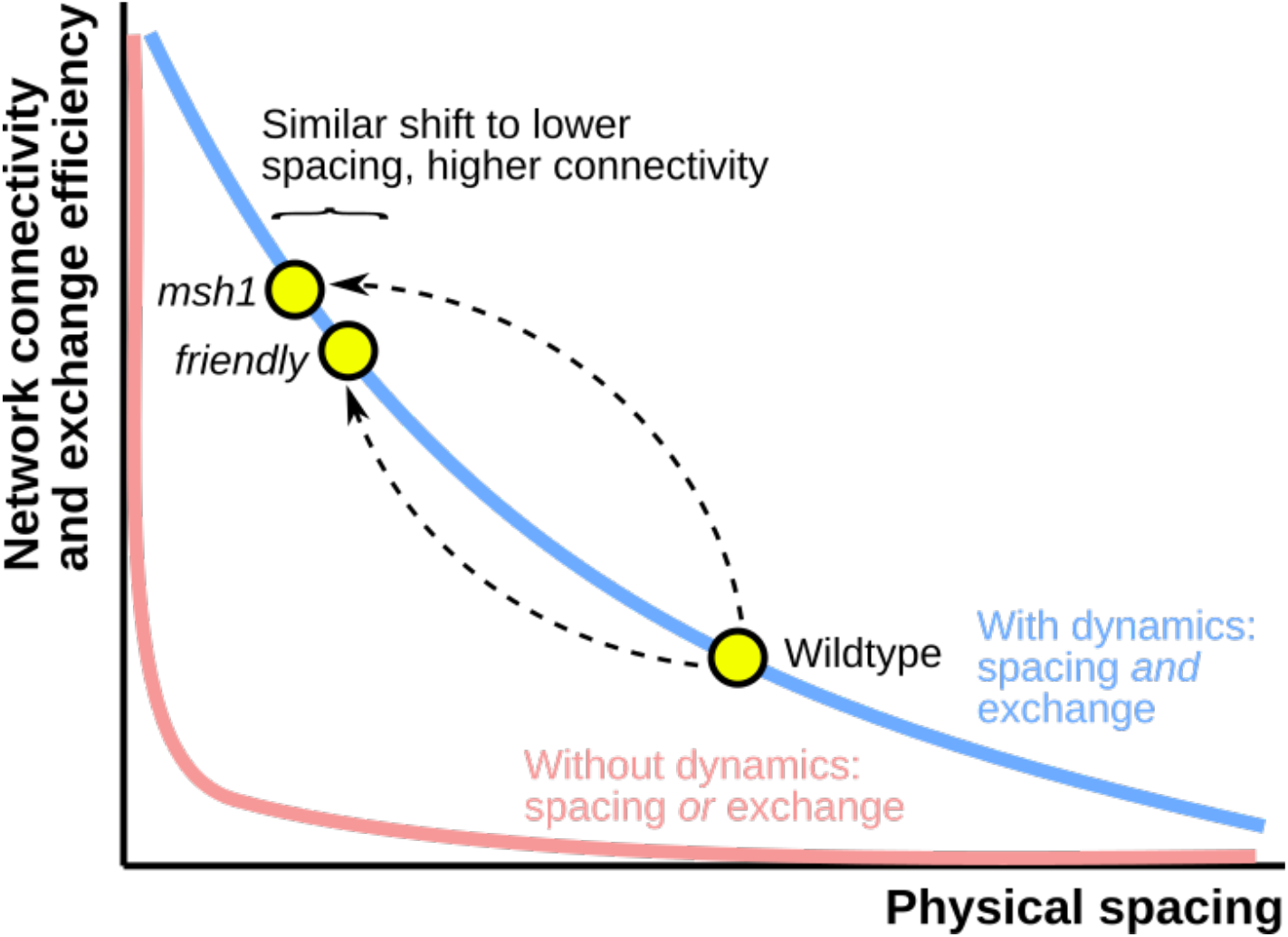
Different resolutions to the social/spacing tradeoff. There exists a tradeoff (coloured curves) between physical spacing of mitochondria (horizontal axis) and the connectivity of the chondriome (vertical axis). Without mitochondrial dynamics, static organelles are either colocalised or spaced, with little capacity to support both behaviours together (pink). Mitochondrial dynamics provides a resolution: as organelles move, they can transiently colocalise while usually remaining spaced (blue), allowing some capacity for both behaviours. Wildtype *Arabidopsis* adopts a particular balance between spacing and encounters. This balance is shifted in strikingly similar ways in the *msh1* mutant compromising organelle DNA maintenance, and the *friendly* mutant compromising mitochondrial dynamics.

The *msh1* mutation has a wide range of influences on the cell. Organelle DNA maintenance is compromised, and many downstream metabolic and structural effects may arise as a result. One potentially quite general principle is that the physical dynamics of organelles exert control on the genetic dynamics of oDNA by dictating which oDNA molecules can interact, be degraded, and so on -- and the cell may thus address genetic priorities by controlling physical behaviour (Johnston, 2019; Edwards *et al*., 2021). Our *msh1* results are not incompatible with the hypothesis that genetic challenges to mtDNA integrity (here, through compromised mtDNA maintenance (Wu *et al*., 2020)) induce a compensatory physical response in mitochondrial dynamics, where the cell sacrifices mitochondrial spacing to allow more encounters. The increased connectivity we observe across the chondriome could then provide individual mitochondria with a chance to access undamaged mtDNA, or extra copies of gene sequences to use as guide strands during double strand break repair. However, the other effects of the *msh1* mutation may also play important or leading roles in shaping the collective dynamic response, including metabolic influence from mitochondrial and chloroplast dysfunction, the further buildup of oDNA mutations, changes to nDNA methylation, and consequent or independent influences on the internal structure of the cell (Davila *et al*., 2011; Christensen, 2014; Wu *et al*., 2020; Xu *et al*., 2011, 2012; Virdi *et al*., 2015; Shao *et al*., 2017). Further work characterising mitochondrial collective dynamics in lines controlling for these influences will help provide further support for the physical-genetic feedback hypothesis.

Mitochondria are increasingly being recognised as ‘social’ organelles, with their interactions playing important functional roles beyond what a collection of independent individuals could achieve (Picard and Sandi, 2020). In plants, a picture of collective behaviour emerging from a population of individuals is particularly pertinent, as mitochondria physically retain individual identities to a much greater extent than in other kingdoms where fused networks are common. The sharing of contents between mitochondria, and consequent control of contents throughout the population, is an example of such emergent behaviour that could not be achieved by independent organelles.

Other examples exist of where plant mitochondrial dynamics may influence mtDNA genetic structure. In plant cells, contrasting with other kingdoms, different mitochondria contain different subsets of the full mtDNA genome (Preuten *et al*., 2010). Many mitochondria may contain no mtDNA at all, while some may contain the full genome (57 genes across 366kb in *Arabidopsis*), and others may contain a subgenomic molecule containing some but not all mtDNA genes (Arimura *et al*., 2004; Gualberto *et al*., 2014; Kozik *et al*., 2019). Processes of mtDNA exchange and recombination are essential to maintain this diverse structure (Bellaoui *et al*., 1998; Arrieta-Montiel *et al*., 2009; Davila *et al*., 2011; Gualberto and Newton, 2017), with mtDNA sharing through the population of mitochondria constituting a ‘discontinuous whole’ (Logan, 2006a). Such sharing and recombination is inherently shaped and limited by the physical behaviour of organelles in the cell (Belliard, Vedel and Pelletier, 1979; Lonsdale *et al*., 1988; Gualberto and Newton, 2017; Aryaman *et al*., 2019; Johnston, 2019; Rose, 2021). In the shoot apical meristem (SAM), a cage-like mitochondrial network has been observed to form (Seguí-Simarro and Staehelin, 2009), in contrast to the largely individual mitochondria observed in other tissues. This network structure allows mtDNA mixing and may facilitate recombination (Edwards *et al*., 2021; Rose, 2021). In conjunction with this physical change, relative expression of MSH1 is particularly high in the SAM, which may both assist with maintenance and support germline mtDNA segregation through gene conversion as an evolutionary priority (Schmid *et al*., 2005; Edwards *et al*., 2021).

These links between the physical behaviour of mitochondria and the genetic behaviour of mtDNA are still being elucidated across kingdoms (Aryaman *et al*., 2019; Johnston, 2019; Edwards *et al*., 2021). The production, degradation, fission, fusion, partitioning, motion, and arrangement of mitochondria in the cell all influence the genetic structure of the mtDNA population. Plant cells, with largely individual mitochondria readily visualised in a quasi-2D cytosolic domain, are an excellent model system for further exploring this link, and we believe that the encounter networks we characterise here will find further use in investigating the vital emergent collective dynamics of the chondriome.

### Experimental Procedures

#### Plant lines

An MSH1 (previously CHM1-1) ethyl methanesulfonate-derived mutant line in the Columbia background generated by G. Redei (Rédei, 1973) was obtained from the *Arabidopsis* stock centre (N3372, http://arabidopsis.info/StockInfo?NASC_id=3372). This line carries an SNP in the fourth exon of genomic region AT3G24320, leading to a nonsynonymous glutamate-> stop codon change. This line was originally isolated in a *gl1* marked plant, a linkage gene in the 3rd chromosome, and so carries a gl1 polymorphism, and lacks trichomes. There is evidence to suggest *gl1* does not alter mitochondrial behaviour (Islam *et al*., 2020), and the gene is highly expressed in only the early SAM, young leaf and young flower, not in the hypocotyl used in this study (Nakabayashi *et al*., 2005; Schmid *et al*., 2005; Klepikova *et al*., 2016). This mutant has been used in previous studies as a disruptor of normal MSH1 function (Xu *et al*., 2011; Wu *et al*., 2020). Seeds of *Arabidopsis thaliana* with mitochondrial-targeted GFP, and the mtGFP-*friendly* (Mito-GFP::*fmt*) line were kindly provided by Prof. David Logan (Logan and Leaver, 2000; El Zawily *et al*., 2014).

### Crossing and DNA extraction

*msh1* and mtGFP seeds were surface sterilized in 50% (v/v) household bleach solution for 4 minutes with continual inversion, rinsed three times with sterile water, and plated onto 1/2 MS Agar. Plated seeds were stratified in the dark for 2 days at 4°C. Seedlings were grown in 16hr light/8hr dark at 21°C for 4-5 days, before transferred to 4:2:1 compost-vermiculite-perlite mixture, and grown until first flower buds developed.

Crossing technique followed the (Browse *et al*., 1993) protocol, with mtGFP plants as the pollen donor and *msh1* accepting. Pollinated stigmas were wrapped gently in plastic wrap and siliques left to develop. F2 seeds were sown onto 50μg/ml Kanamycin 1/2 MS plates, selecting for individuals carrying the fluorescence construct (Logan and Leaver, 2000), and grown on soil as before. Leaf samples were taken for DNA extraction from all but F2 seeds.

Quick DNA extraction was performed on young leaf samples (2-3 weeks old, age dependent on growth rate). Leaf samples were macerated with a pipette tip in 40μl Extraction Buffer (2.5mL 2M TRIS-HCL, 500μL 1M EDTA, 6.25mL 2M KCL, made to 50mL with BPC water).

Sample was then incubated in a heat block for 10min at 95°C. 40μl dilution buffer was then added (3% BSA (1.5g in 50mL), filter sterilised), and samples spun down at 13000rpm for 60s before storing at -20°C.

### Genotyping and sequencing

For genotyping, primer set 1 was used. A reverse primer (RP1) running into the snp site was designed using dCAPS finder 2.0 (Neff *et al*., 1998), and the forward primer (FP1, see supplementary material) was designed 200bp upstream of the restriction site. By design, BsrGI will cut a region of 30bp from the 293bp element if the SNP is present, producing one larger (260bp) and one smaller (∼30bp) fragment compared to the WT single fragment (293bp). After PCR amplification, half (5μL) of PCR product for each sample was directly added to 1.5μL Cutsmart buffer [NEB], 0.2μL BsrGI restriction enzyme [NEB], 8.3μL nuclease-free H_2_O. Samples were then incubated at 37°C overnight, before alternate undigested/digested samples loaded for gel electrophoresis.

To sequence *MSH1*, the region of interest was first amplified by PCR using primer set 2 (see supplementary information) and Phusion high-fidelity DNA polymerase (NEB CAT#M0530S). PCR products were then purified using QIAquick PCR Purification Kit (Qiagen) and sequenced from primer FP2 using an ABI 3730 capillary sequencer (Applied Biosystems).

### Imaging and video analysis

Seedlings for imaging were sterilized, stratified and grown 50μg/ml Kanamycin 1/2 MS plates as described above. After 4-5 days, seedlings were taken for imaging, and prior to mounting, stained with 10μM propidium iodide (PI) solution for three minutes to capture the cell wall. Simple mounting of whole seedlings on microscope slides with coverslips was used (modified from (Whelan and Murcha, 2015)). In order to minimize the effects of hypoxia and physical stress on the seedling, imaging was undertaken in less than ten minutes after the cover slip was added.

We used a Zeiss 710 laser scanning confocal microscope for imaging of seedlings. To characterise cells we used excitation wavelength 543nm, detection range 578-718nm for both chlorophyll autofluorescence (peak emission 679.5nm) and for PI (peak emission 648nm). For mitochondrial capture we used excitation wavelength 488nm, detection range 494-578nm for GFP (peak emission 535.5nm). Time-lapse images were taken, and all samples used in this study have the same time interval between frames, and same length of capture, allowing for direct comparison.

For image analysis, single cells were cropped using the PI cell wall outline with Fiji (Image J 2.0.0). The universal length scale of 5 pixels/μm was applied across all samples. To counter the occasional sample drift within time-lapse videos, 3D drift correction was applied with default settings, using the cell outline via the Propidium Iodide channel as the stability landmark (correct 3D drift, FIJI, ImageJ 2.1.0, (Parslow, Cardona and Bryson-Richardson, 2014)).

Tracking of individual mitochondria was done using Trackmate (Tinevez *et al*., 2017) in ImageJ 2.0.0. The LoG detector was used with typical settings being 1μm blob diameters (the typical size of a mitochondrion), although 0.8μm was occasionally used for lower signal samples. Detection threshold was set between 1.5-8, and filters applied on spots if necessary. The Simple LAP Tracker was run with a linking max distance of 4μm (3μm used for a few samples), gap-closing distance of 5μm (4μm used for a few samples) and gap-closing max frame gap of 2 frames. For each sample, quality of overlaying detection for mitochondria was scrutinised, and occasional tracks edited for precision.

### Physical statistics

Physical statistics include speed (μm/frame), the distance moved per frame per trajectory. This value is averaged over all trajectories from the duration of the video. Inter-mitochondrial distance is the minimum Euclidean distance (μm) between every mitochondrion and its nearest physical neighbour in each frame. This value is average over all frames of the video. Colocalisation time is the number of frames any two mitochondria have spent within a threshold distance (1.6μm) of each other, averaged over all frames.

### Network statistics

Encounter networks are built from the close associations of mitochondria. A threshold distance of 1.6μm was used to define a characteristic close association, being just over one mitochondrion’s length. Lower threshold distances can also be used, yielding less encounters, but similar connectivity trends (Chustecki *et al*., 2021). Networks build up as encounters (edges) between mitochondria (nodes) are registered over time.

The mean degree is the number of immediate neighbours each node has, averaged over the number of nodes in the network. Network efficiency is the average, over all pairs of nodes, of the reciprocal shortest distance between each pair:

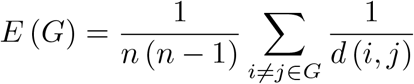

where G is the network of interest, *n* is the number of nodes in the network and *d*(*i, j*) is the distance (edge number) between node *i* and node *j*. The graph diameter is the length of the longest direct path across the network, a quantification of the number of edges connecting the two furthest nodes across a network. The mean graph betweenness centrality is the average number of shortest paths crossing each node in the network. The mean connected component number is the average number of disconnected subgraphs within the network.

### Accession numbers

All analysis code and data is available from Github at https://github.com/StochasticBiology/plant-mito-dynamics

## Acknowledgments

J.M.C. is supported by the BBSRC and University of Birmingham via the MIBTP doctoral training scheme (grant number BB/M01116X/1). This project has received funding from the European Research Council (ERC) under the European Union’s Horizon 2020 research and innovation programme (grant agreement nos. 805046 (EvoConBiO) to I.G.J. and 715441 (GasPlaNt) to D.J.G., and supporting R.D.E).

We gratefully acknowledge the Imaging Suite (BALM) at the University of Birmingham for support of imaging experiments and thank Alessandro di Maio and Prof. Markus Schwarzländer for advice with design and analysis of imaging experiments. We are very grateful to Prof. David Logan for mtGFP seeds. Thanks to Miguel Pachón Peñalba and Alice Darbyshire for molecular biology experimental advice and help in primer design. We gratefully acknowledge the services of Genomics at Birmingham for sequencing experiments.

## Author Contributions

IGJ conceived the project. JMC created plant lines. JMC and RDE performed sequencing and validation. JMC performed microscopy. JMC performed statistical analysis. IGJ and DJG supervised laboratory work. IGJ supervised theoretical work. IGJ and JMC drafted the manuscript. All authors edited the manuscript.

## Legends for Supporting Information

**Supplementary Figure 1: Genotyping for F3 *msh1* homozygosity leads to consistently variegated F4 progeny.**

**Supplementary Figure 2: Plant phenotypes reveal developmental differences across genotypes.**

**Supplementary Figure 3: Single nucleotide polymorphism in MSH1 retained in the F3 generation of mtGFP-*msh1* cross.**

**Supplementary Figure 4: Sample encounter networks for mtGFP and mtGFP-*msh1*. Supplementary Figure 5: No evidence found for a difference between median cell area across genotypes.**

**Supplementary Figure 6: Node number and edge number of encounter networks did not vary greatly between lines for mtGFP, mtGFP-*msh1*, and mtGFP-*friendly*.**

**Supplementary Figure 7: Social summary statistics provide evidence of differences between mtGFP, mtGFP-*msh1* and *friendly*, at three earlier time points.**

**Supplementary Video 1: An example cell from 4-5 day old mtGFP-*msh1* hypocotyl, showing GFP-tagged mitochondria**.

## Supplementary Information

**Supplementary Figure 1:**
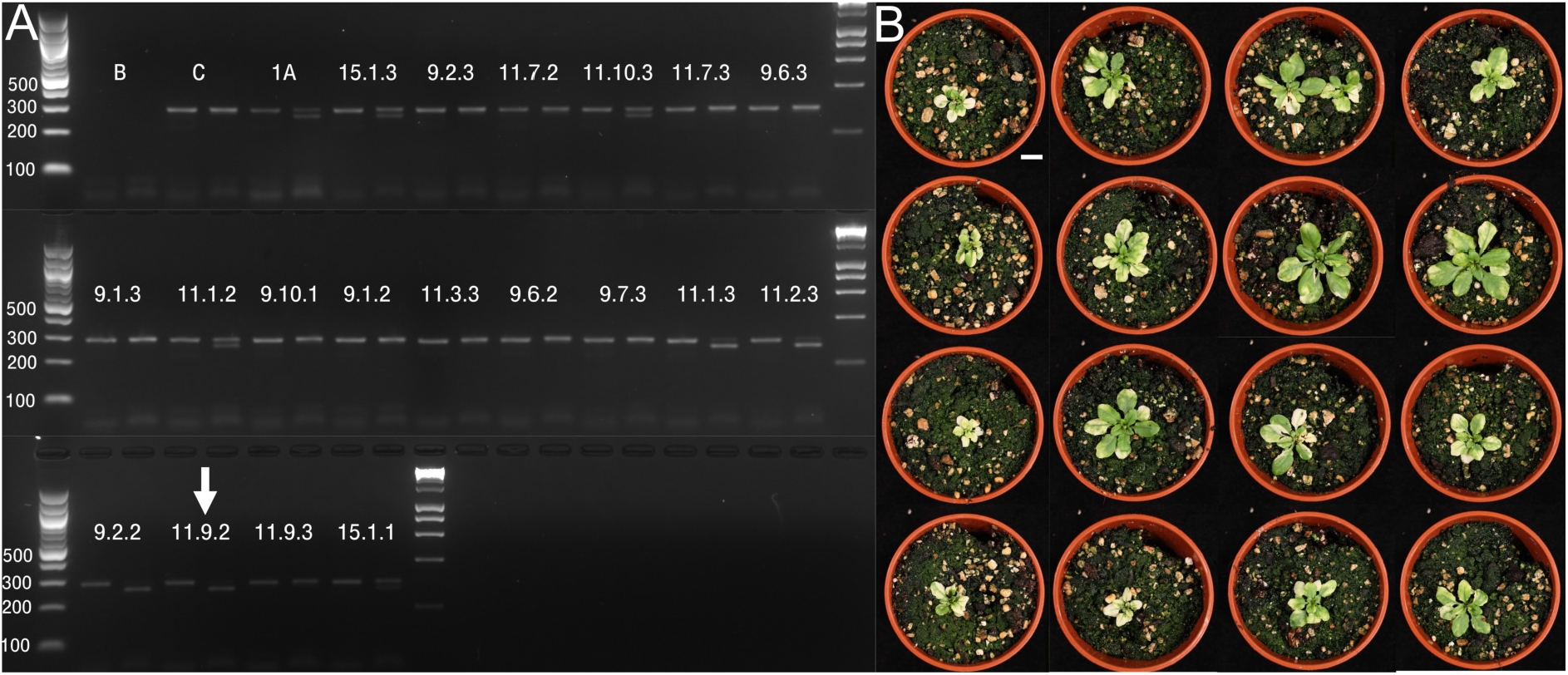
Genotyping for F3 *msh1* homozygosity leads to consistently variegated F4 progeny. (A) dCAPS genotyping for WT gives 293bp fragment, but in the presence of *msh1* SNP mutation gives ∼260bp fragment, when digested with a restriction enzyme. Each line has an undigested (left band) and a digested (right band) sample. Homozygosity is demonstrated by one upper band (left, 293bp), and one lower band (right, ∼260bp). Heterozygosity is demonstrated by one left band and two fragments in the right band. (B) Phenotype of candidate line 11.9.2, showing all individuals with variegated phenotype typical of the *msh1* mutation in *Arabidopsis* (30 days old). Scale bar = 1cm.

**Supplementary Figure 2:**
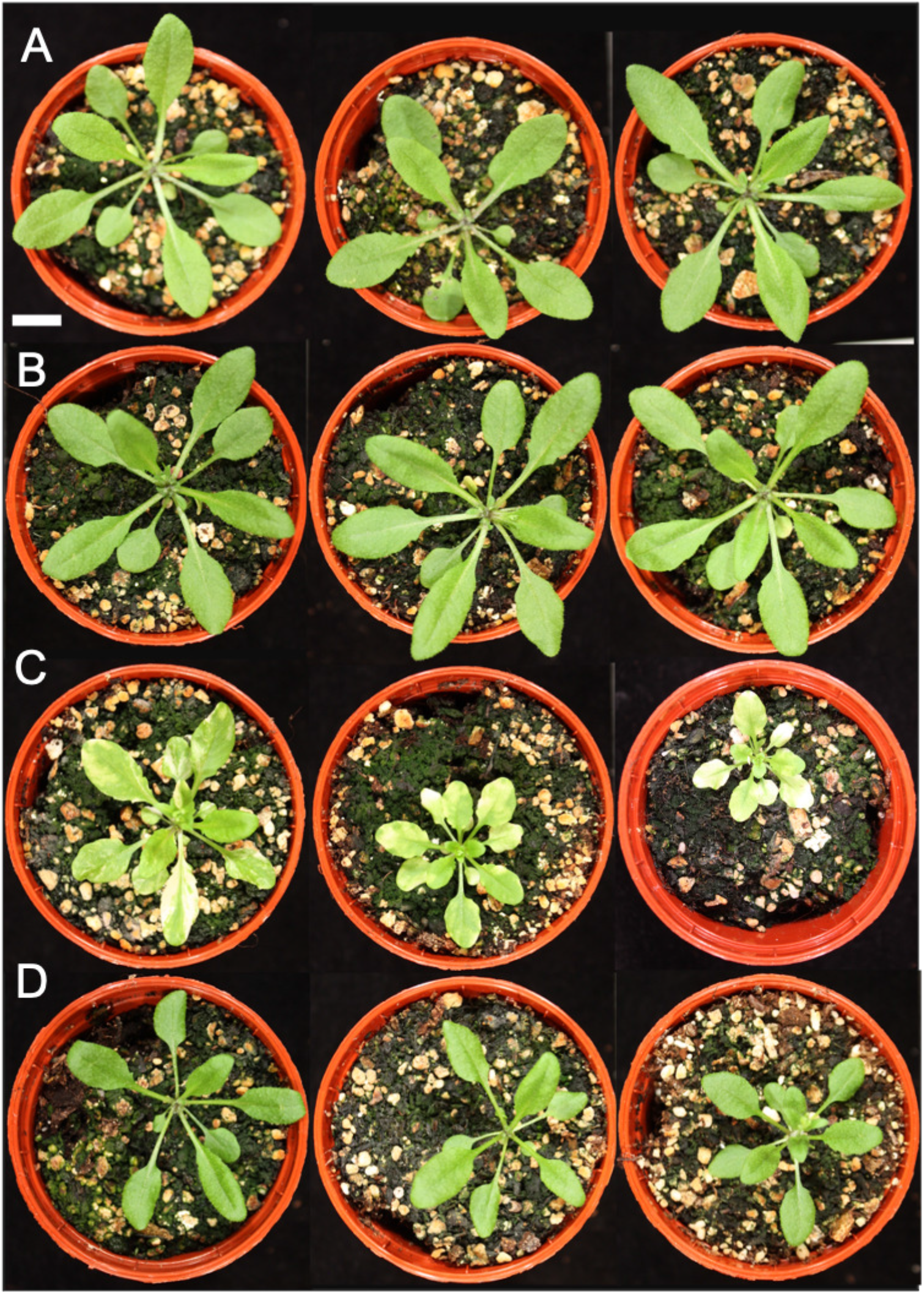
Plant phenotypes reveal developmental differences across genotypes. Rosette images of 31 day old plants taken of A) Col-0; B) mtGFP; C) mtGFP-*msh1*; D) mtGFP-*friendly*. Scale bar = 1cm.

**Supplementary Figure 3:**
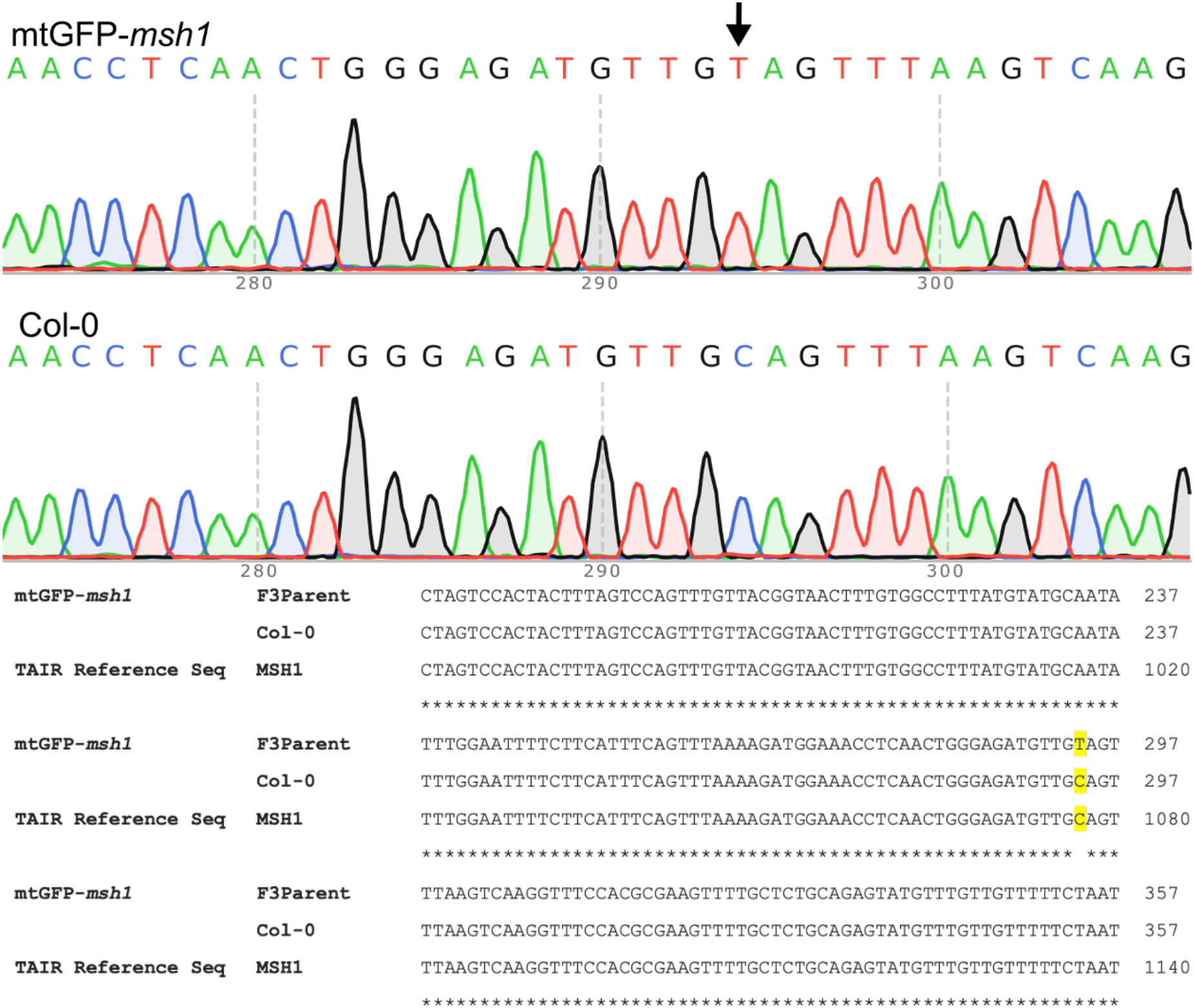
Single nucleotide polymorphism in MSH1 retained in the F3 generation of mtGFP-*msh1* cross. Upper panel illustrates single autoscaled peaks, showing base pair reads across the middle of the amplified region and at position 294 (arrow), evidence of a homozygous SNP. Lower panel shows alignment of base pair reads of mtGFP-*msh1* F3 parent, Col-0 sample, and the TAIR reference genome at the *MSH1* gene. Highlighted base shows the SNP leading to CAG (glutamine) to TAG (stop).

**Supplementary Figure 4:**
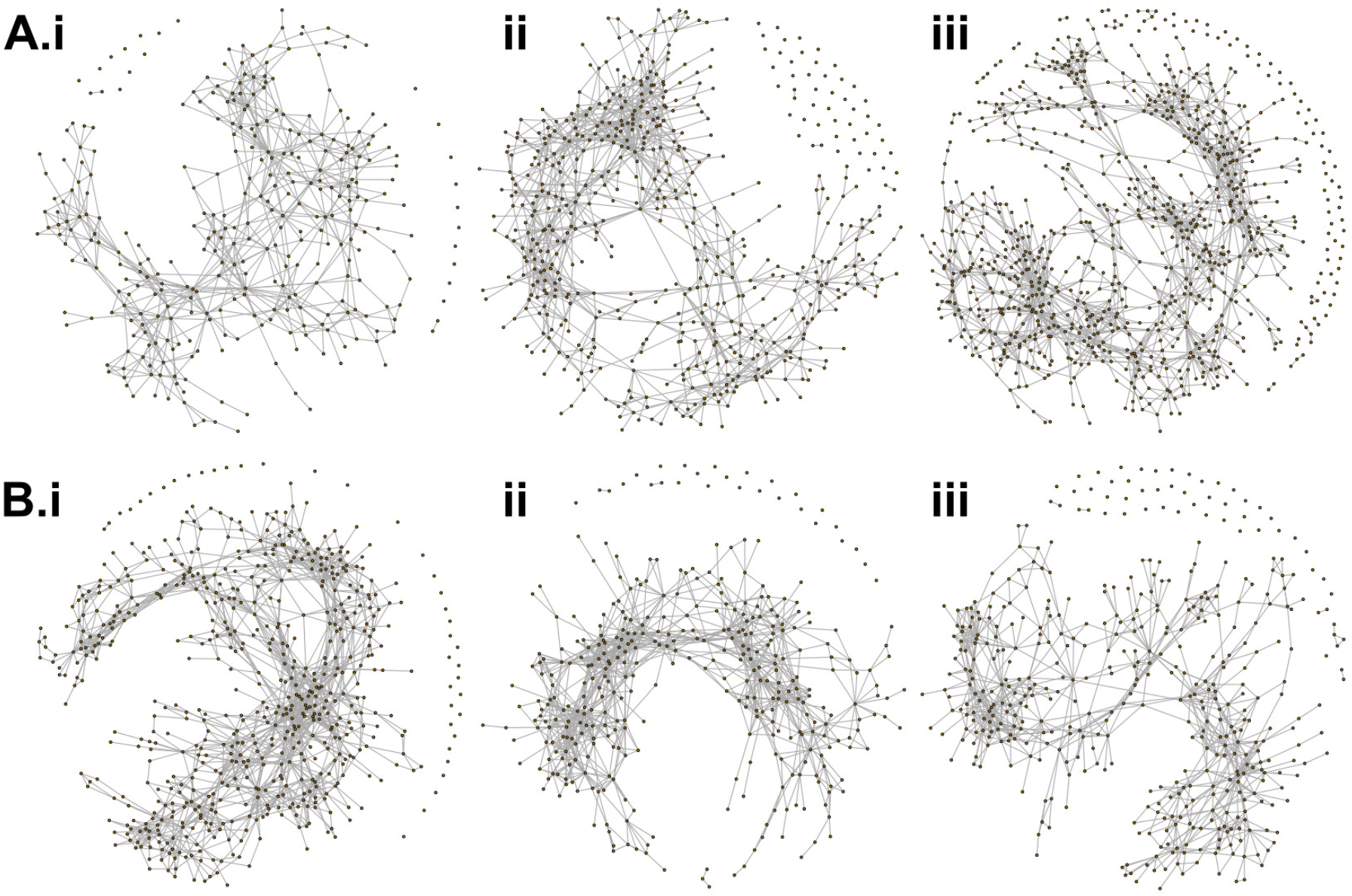
Sample encounter networks for mtGFP (A) and mtGFP-*msh1* (B). Networks are built from close encounters (edges) between mitochondria (nodes) (see methods) over different cells (i)-(iii). Networks here are built up from 233 seconds of video time.

**Supplementary Figure 5:**
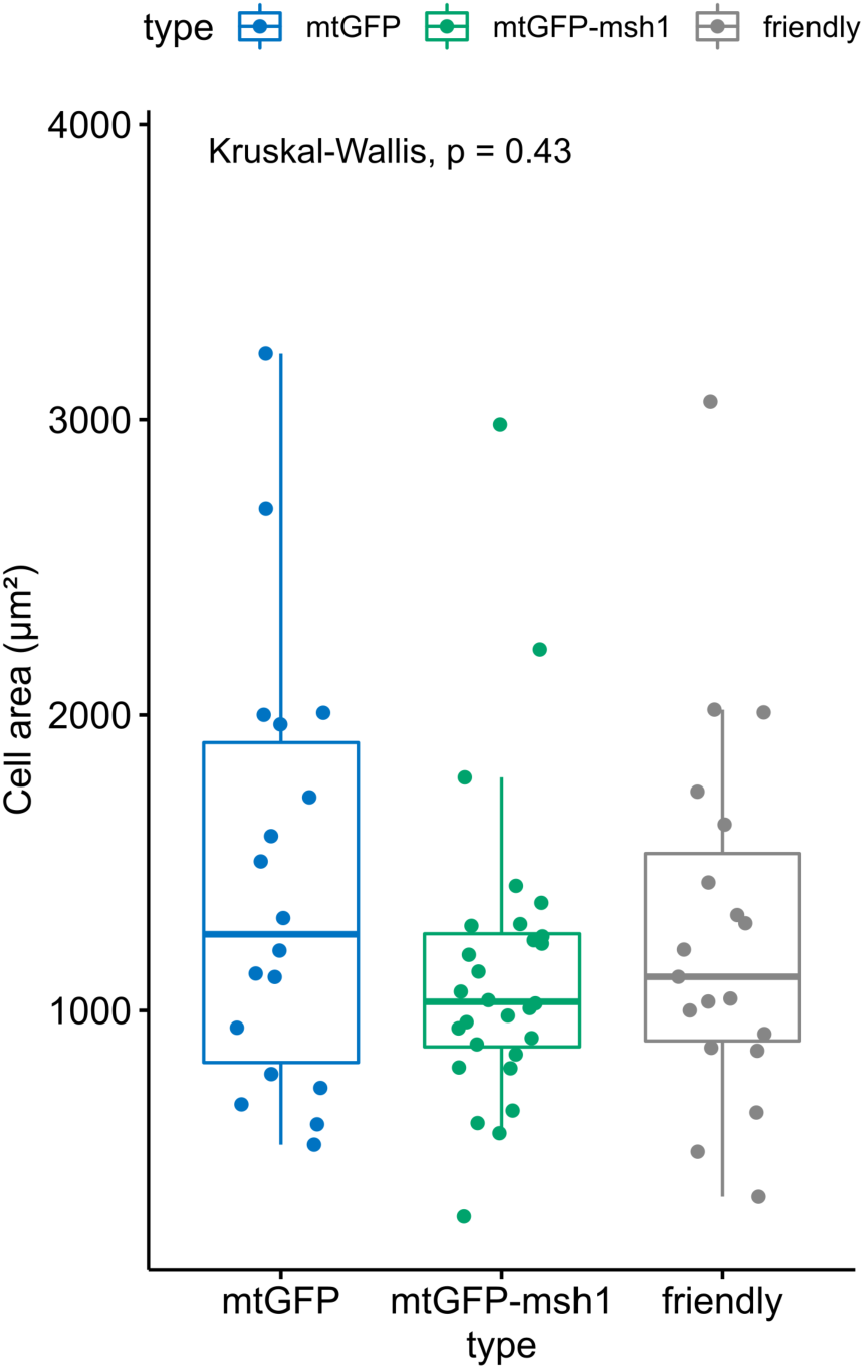
No evidence found for a difference between median cell area across genotypes. Comparison of two-dimensional cell area (μm^2^) between the three genotypes using the Kruskal-Wallis test. Boxplots represent the median and 25th/75th percentile, with whiskers showing the smallest/largest value within 1.5x the interquartile range. P-value represents Kruskal Wallis test outcome across all three genotypes.

**Supplementary Figure 6:**
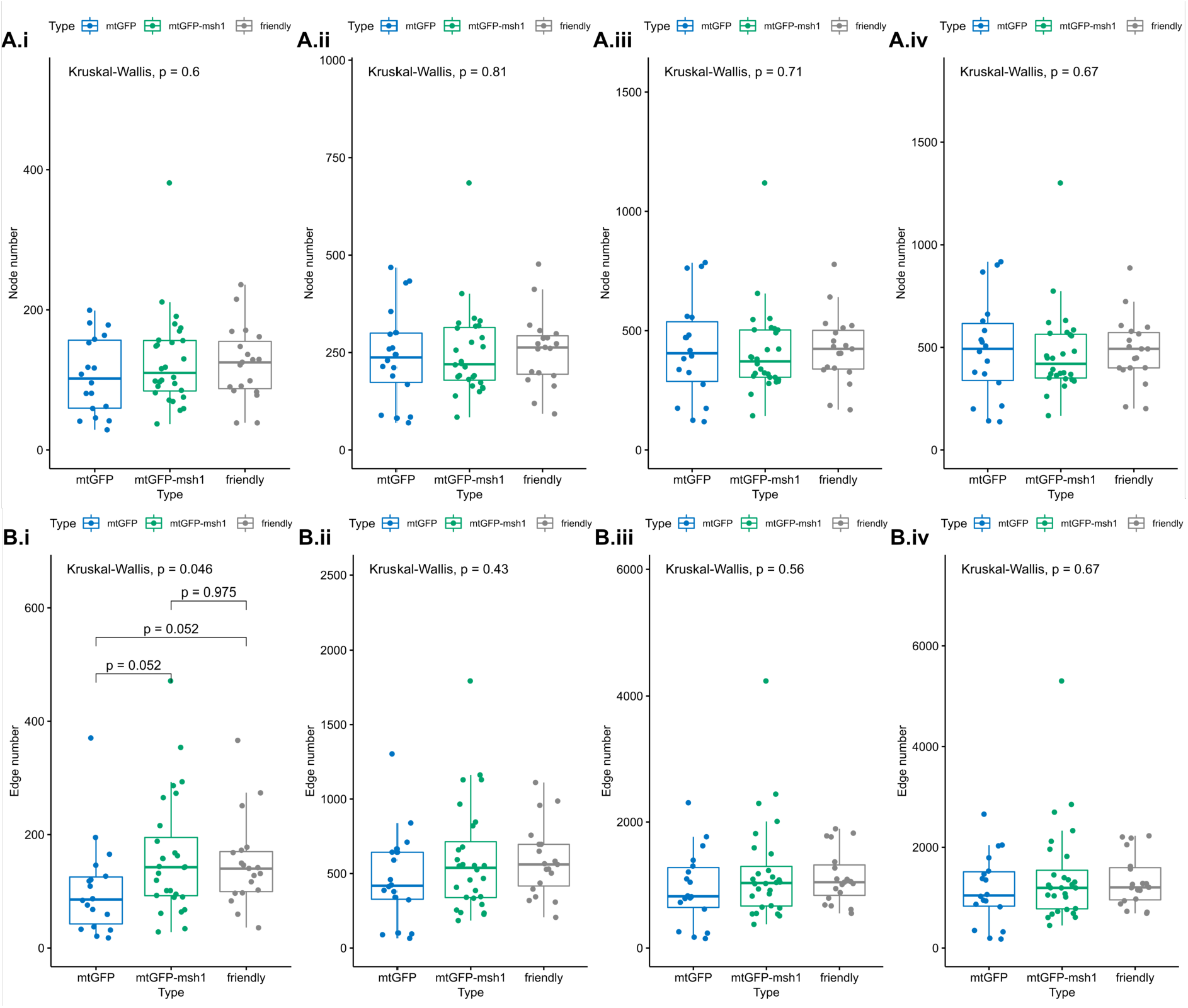
Node number (A) and edge number (B) of encounter networks did not vary substantially between lines for mtGFP, mtGFP-*msh1*, and mtGFP*-friendly*. With the exception of edge numbers for early frame 10. From left to right (i-iv) graphs show snapshots of networks at frames 10, 50, 100, 120. P-values represent Kruskal Wallis test outcomes across all three genotypes, and pairwise p-values are false discovery rate adjusted outcomes of a post-hoc Dunn test, without multiple hypothesis correction across statistics.

**Supplementary Figure 7:**
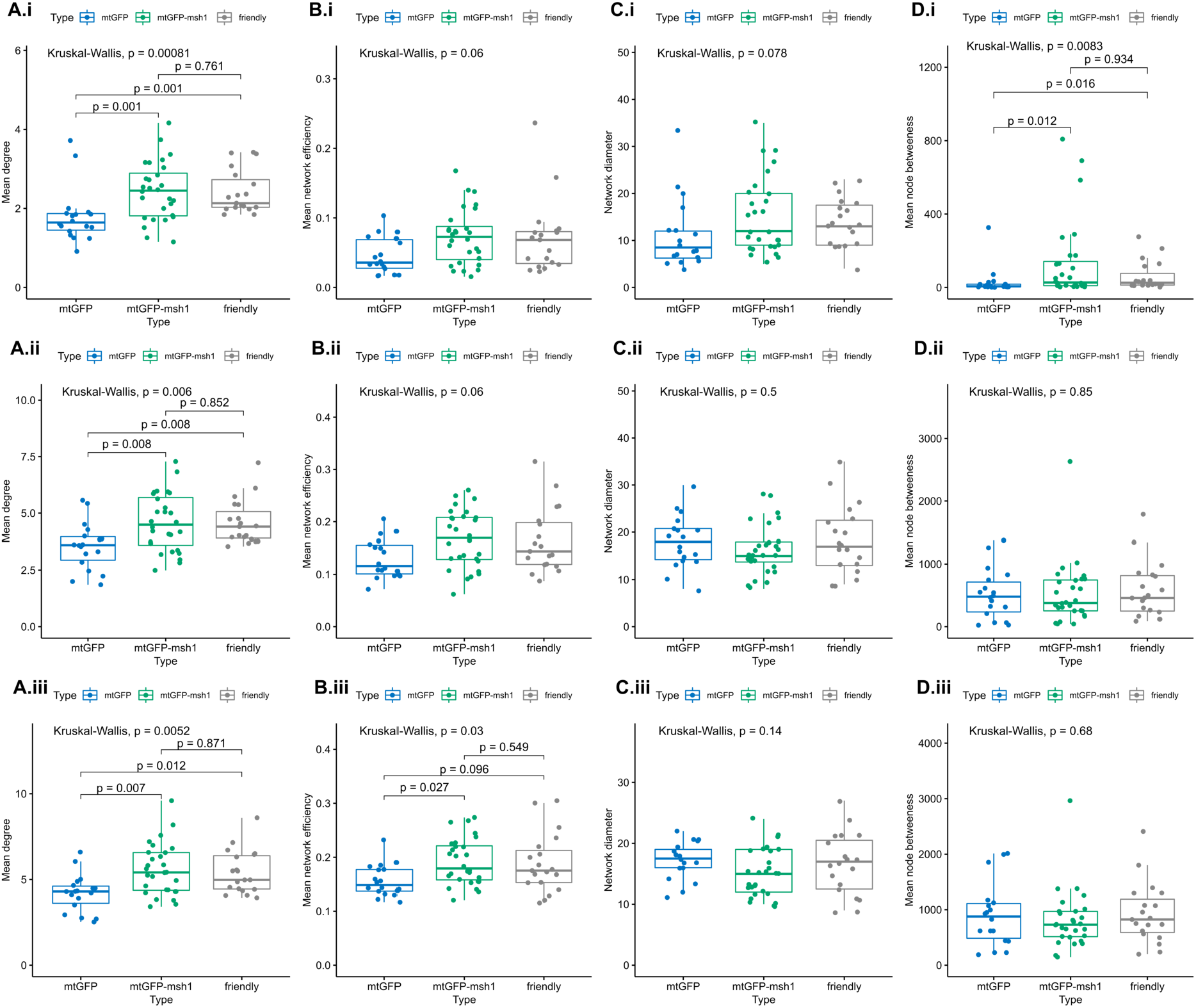
Social summary statistics (A-D) provide evidence of differences between mtGFP, mtGFP-*msh1* and *friendly*, at three earlier time points (10 frames (i), 50 frames (ii), 100 frames (iii). Each point represents a summary statistic for one cell (mtGFP *n*= 18, mtGFP-*msh1 n*= 28, *friendly n* = 19). P-values represent Kruskal Wallis test outcomes across all three genotypes, and pairwise p-values are false discovery rare adjusted outcomes of a post-hoc Dunn test, without multiple hypothesis correction across statistics. Boxplots represent the median and 25th/75th percentile, with whiskers showing the smallest/largest value within 1.5x the interquartile range. Frames correspond to 19, 97 and 194 seconds, respectively. P-values are for individual experiments.

Link: https://org.uib.no/stochasticbiology/SuppVideo1-MSH17.avi

**Supplementary Video 1: An example cell from 4-5 day old mtGFP-*msh1* hypocotyl, showing GFP-tagged mitochondria** (green), and a Propidium Iodide stain around the cell (red); autofluorescence from the chloroplasts also detected (red).

